# Dynamic changes of protist community composition along a surface water-groundwater transect in the Danube wetland Lobau, Vienna, Austria

**DOI:** 10.1101/2025.07.30.667637

**Authors:** Angela Cukusic, Clemens Karwautz, Christina Murhammer, Petra Pjevac, Grit Rasch, Christine Stumpp, Christian Griebler

## Abstract

Groundwater remains an unexplored habitat in terms of protistan diversity and community dynamics, when compared to better-studied surface aquatic environments. To address this knowledge gap, a comparison of protistan communities found at two differing sites, a wetland surface water body and the close-by shallow aquifer, aims to shed light on environmental drivers of both the total and active protist communities using molecular analyses. The two-year time series with monthly samplings provided insight into seasonal pattern, and barcoding allowed taxonomic affiliation. Our study revealed clear differences in community composition and trophic modes with the habitat type. Protistan communities in shallow groundwater exhibited pronounced seasonal dynamics somehow temporally linked to the surface water counterparts. Higher absolute water levels in the backwater compared to groundwater and the significant fraction of phototrophic protists sampled from the shallow aquifer point at groundwater recharge from surface water to strongly influence protistan community composition.

## 1 INTRODUCTION

Investigations of microbial communities in transition zones are critical to our understanding of community assembly, adaptations to changing environmental conditions, ecosystem processes, and diversity of life on Earth. The importance of surface–subsurface exchanges as part of the hydrological cycle, for biogeochemical processes and biodiversity has repeatedly been addressed in the past (Malard et al., 2002; Hou et al., 2017), and we know that groundwater-dependent ecosystems such as wetlands perform critical watershed functions that affect the spatio-temporal dynamics and thus the rates of material and energy fluxes (Lane et al., 2023; Ward et al., 1999). Water moving between surface waters and groundwater triggers and underlies changes in physico-chemical conditions and actively transports microorganisms from one place of specific selection pressures (selection, drift, speciation and dispersion Hanson et al. (2012)) to another. This movement and transport of unicellular organisms between terrestrial soil and aquatic habitats seems to occur frequently (Sieber et al., 2020), and with routes of passive dispersal to subsurface via groundwater recharge from seepage and surface waters.

Aquatic protists, a heterogeneous group of microeukaryotes, are characterized by complex phylogenetic relationships, diverse trophic modes and a wide spectrum of ecological roles, they contribute to key biogeochemical cycles and connect the microbial world with aquatic and terrestrial metazoan communities (Gadd and Raven, 2010; Azam et al., 1983). In comparison to Bacteria and Archaea, protists have a more restricted range of metabolic capabilities encoded in their genome, however ranging from primary production and mixotrophy to grazing of prokaryotic cells and virus particles and parasitic lifestyle (Sherr and Sherr, 1988; Dodds and Whiles, 2010; Falkowski et al., 2008). Their functional potential and niche space are more defined than for prokaryotes. Yet, ecological studies of protist communities are often limited to a few environments (e.g. marine environments or surface inland waters) (Bik et al., 2012). Research on groundwater protists is rare and partly outdated, largely based on morphological identification (Sinclair and Ghiorse, 1987; Hirsch and Rades-Rohkohl, 1983), quantitative data from microscopic counts (Novarino et al., 1997; Karwautz et al., 2022), and molecular analysis using fingerprinting methods (Euringer and Lueders, 2008). Most studies largely focused either on contamination plumes or peculiar ecosystems not representative of general groundwater properties (Kinner et al., 2002; Novarino et al., 1994; Holmes et al., 2013; Oliverio et al., 2018; Borgonie et al., 2015). However, the notable exceptions that combined cultivation-based approaches and quantitative assessments with low-throughput sequencing have substantially advanced our understanding of groundwater protistan communities (Risse-Buhl et al., 2013; Loquay et al., 2009).

The usage of new high-throughput sequencing technologies has recently emerged as a suitable method for identifying protist contributions to the groundwater food web (Herrmann et al., 2020), but little progress has been made to link the distribution to the driving environmental conditions. The 18S rDNA barcoding as a means to link environmental and spatial drivers of protist communities has so far been used predominantly for soils (Oliverio et al., 2020; Huang et al., 2023), marine (Egge et al., 2021; Caracciolo et al., 2022), and surface freshwater ecosystems (Olefeld et al., 2020; Lepère et al., 2013), largely leaving out groundwater, with even the most recent review in the field entirely neglects groundwater ecosystems (Singer et al., 2021).

In this study the protist community composition and diversity of an oxbow lake is directly compared to the community in shallow groundwater 25 meters distant from the shoreline. The shallow aquifer differs from the surface water in many ways besides the obvious lack of light, its trophic state, oxygen availability, the amplitude of seasonal environmental dynamics and thus the occurring organisms. Groundwater poses specific challenges as a habitat, and is typically inhabited by well-adapted organisms assembling to diverse communities, distinct from those found at the surface. However, the consequences of hydrological exchange between the surface water and the shallow aquifer to dynamics in protistan community composition remain unclear, particularly in regards to the proportion of generalist taxa, i.e. those consistently detected in both environments, relative to specialists, restricted to either surface or subsurface habitats. In addition to passive dispersal, protists are capable of actively moving both by gliding and swimming, and some are also capable of stably attaching to surfaces. Taxa associated to sediment matrices are likely to be attached or gliding, such as cercozoans and amoeba, whereas the free swimming taxa and filter-feeding flagellates are hypothesized to be more abundant in the water column of aquatic habitats. Still, their feeding mechanisms could largely be similar, and include interception feeding, filter feeding and grazing.

Based on morphological identification, the biggest portion of groundwater protist communities is made up of heterotrophic flagellates, such as bodonids, cercomonads and cryptomonads, with their absolute numbers depending on organic load and availability (Novarino et al., 1997; Risse-Buhl et al., 2013; Loquay et al., 2009). First instances of molecular barcoding put ciliates as most abundant taxa in karstic aquifers (Herrmann et al., 2020), but the ciliate bias caused by the characteristic multiple ribosomal transcription is still an issue to be addressed in protistan barcoding (Medinger et al., 2010). Other than organic load, it is difficult to discern which of the environmental filters that are known to shape surface communities are of importance in groundwater. The phototrophic taxa which play a significant role in diversity and activity of surface habitats are not expected to prosper in groundwater, and could make a good indicator for surface water infiltration. The pH and mean annual precipitation trends that were shown to influence soil communities (Oliverio et al., 2020), are secondary to site-specificity and water quality effect when it comes to wetlands (An et al., 2023), and yet seasonal drivers and a clear yearly cycle was recently studied in brooks and ponds(Simon et al., 2015), and in lakes (Cruaud et al., 2019), where wet and dry season defined community composition. This goes along with the well-established seasonal fluctuations of freshwater diatoms and dinoflagellates (Sommer et al., 1986). The protist seasonality, so pronounced in most ecosystems, might be expected to an extent even in groundwater, especially in shallow aquifers, but unlikely being the most important driving factor. Anthropogenic influences were present in some cases (Garner et al., 2022), as were the impacts of climate change (Gerasimova and et al., 2024), further showing the need for establishing the groundwater protist baseline before it starts changing in the anthropocene.

The aim of this study was to identify the abundant protist taxa in a backwater and neighbouring shallow groundwater in the wetlands of the Danube Floodplain National Park, in Austria. Samples were collected monthly in the span of two years, a period which has previously been proven sufficient for capturing protist seasonality trends in inland waters (Cruaud et al., 2019; Simon et al., 2015). High-troughput sequencing of the V9 region at the end of the 18S rDNA gene was applied with previously tested primers (Geisen et al., 2019) to allow uncovering protist community composition and to compare against results of other 18S-based studies, specifically those from groundwater ecosystems (Herrmann et al., 2020).

We hypothesized: **(i)** a clear distinction between the environmental conditions in surface water and groundwater accompanied by a pronounced difference in the protistan community composition of the two, **(ii)** groundwater protist diversity lower than in the surface water, **(iii)** a dominance of non-photoautotrophic taxa in groundwater, as a result of the specific environmental conditions, and **(iv)** seasonal dynamics being more pronounced in surface water. Furthermore, considering that the Lobau groundwater sampling site is likely to be recharged at a higher rate in winter months (Dreher et al., 2006), **(v)** a higher similarity of groundwater protistan communities to the surface water communities was hypothesised for the winter season.

## 2 MATERIALS AND METHODS

### 2.1 Sampling and study area and sampling

From summer 2020 to summer 2022 sampling of surface water and groundwater was performed in the area of the Viennese Danube wetlands, Austria, located in the national park Lobau. Three groundwater wells and one backwater (Fig. 1) were sampled on a monthly basis, but only the water samples from the 10 meter deep groundwater well D15 and the backwater Eberschüttwasser (ESW) were processed for protists community analysis, representing the far ends in environmental conditions between the surface water and the groundwater.

**Figure 1.**
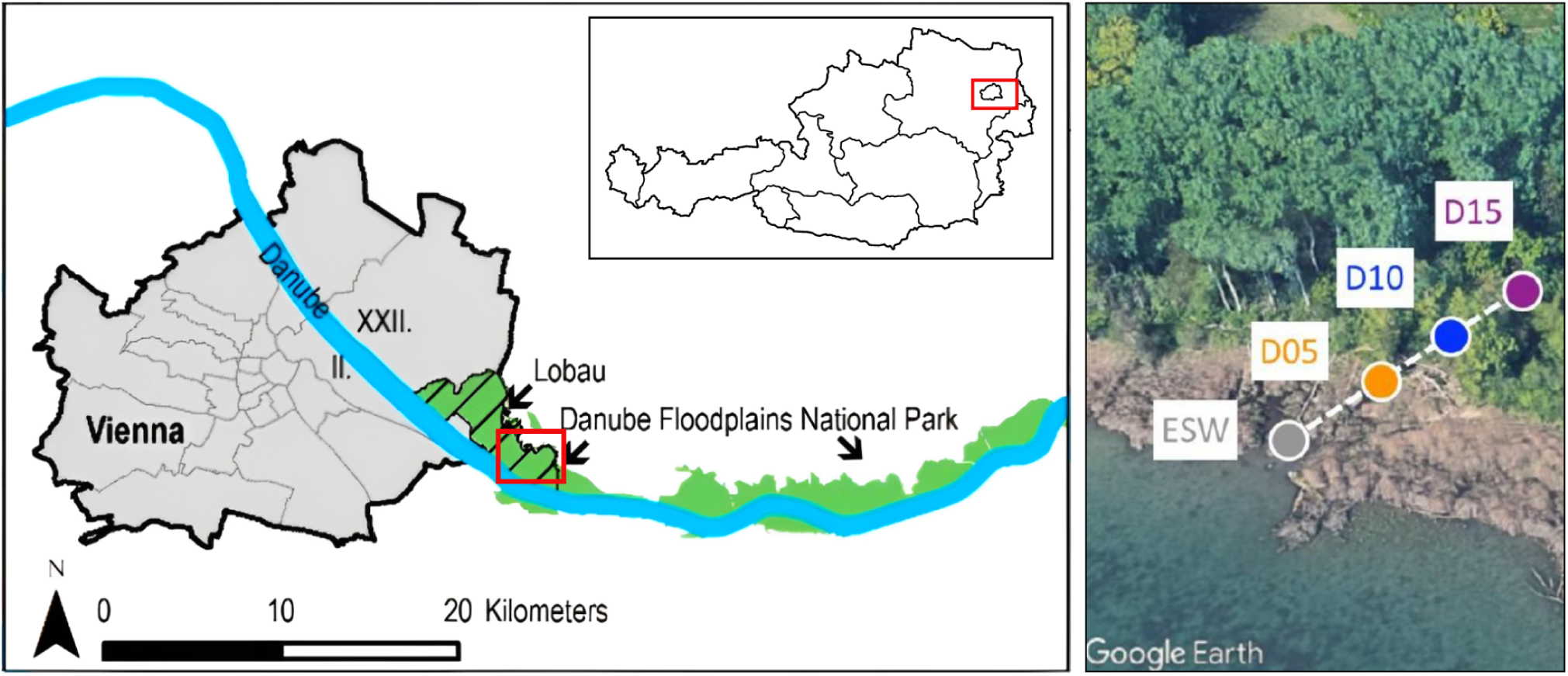
**Left**. Location of the sampling sites in Vienna, Austria. Source: adapted from (Arnberger et al., 2021). **Right**. Positioning of the Lobau sampling sites. Sites D15 and ESW were analysed for protist communites.

Groundwater was pumped with a submersible pump that was submerged to 6 meter depth, for two well volumes, in attempts to remove the stagnant water in the well. After pre-pumping and stability in key environmental variables, temperature, oxygen, electrical conductivity and pH were recorded by means of field sensors (WTW Multi 3620, Weilheim, Germany). Groundwater samples for hydrochemical analyses were filled without any treatment into 50 mL plastic tubes. Water samples for DIC analysis were filtered on-site through a 0.45 *µ*m syringe filter. The same was done for DOC samples with an additional step of adding HCl to reach a pH*<*2. Water samples intended for molecular analysis were collected into 10 L carboys and filtered back in the lab through 0.22 *µ*m Sterivex filter. Filters were stored at −20 °C until they could be further processed.

### 2.2 Physico-chemical and microbial parameters measurement

Concentrations of major ions (Na^+^, K^+^, Ca^2+^, Mg^2+^, Cl^-^, SO_4_^2-^) were measured by ion chromatography (Dionex ICS1100 RFIC, Thermo Scientific, Idstein, Germany). The green house gases carbon dioxide (CO_2_) and methane (CH_4_) were measured via laser spectroscopy (Picarro, G2201-i). Dissolved organic and inorganic carbon (DOC and DIC, respectively) were quantified with a TOC-L analyser (Shimadzu TOC-L). The microbial activity (total and external ATP) was measured based on the Hammes et al. (2010) protocol, with the same adjustments as in Retter et al. (2023). It was performed with the BacTiter-Glo Assay (Promega, USA) in a luminator plate reader (Promega, Madison, WI, USA). Internal activity, used in the analyses and refered to as the ATP, was calculated as the difference between the total and external ATP. The total counts of prokaryotic cells (TCC) were measured with a Cytek® Amnis® CellStream® flow cytometer (Amnis CellStream, Luminex, Austin, TX, USA), after being stained with nucleic acid dye SYBR Green I (Invitrogen, Darmstadt, Germany), using the same settings as Retter et al. (2023).

### 2.3 DNA and RNA extraction, Illumina sequencing and sequence processing

The genetic material was extracted based on a phenol-chloroform extraction protocol optimised for the co-extraction of RNA and DNA (Lueders et al., 2004) with some modification (Lever et al., 2015) to prepare for the 18S rDNA gene amplicon sequencing. The 0.22 *µ*m Sterivex filters which contained the microbial biomass were cut into small pieces and put into a 2 ml tubes containing zirconia-silica beads (Biospec, Bartlesville, USA), in a 1:1 ratio of 0.1-mm and 0.7-mm size. Next, 160 mol/L NaPO_4_, 3.191×10-5 mol/L TNS and 11.4 mol/L PCI (phenol-chloroform isoamyl alcohol in 25:24:1) were added and cells were lysed via beadbeating (FastPrep24, MP) for 45 seconds at 6.5 m/s. The samples were centrifuged (13 000 g and 4 °C) for 4 minutes, and the resulting supernatant was taken (900 *µ*l) and extracted with 1 volume PCI (11.4 mol/L). After centrifugation, 800 *µ*L of the supernatant was mixed with 0.011 mol/L CI (24:1 chloroform:isoamyl alcohol) and transferred to a “Phase Lock Gel Light” column (Eppendorf, Hamburg, Germany), separating (13 000 rpm, 4 °C, 4 mins) the nucleic acids into the upper aqueous phase. The supernatant (650 *µ*l) was mixed with two volumes PEG buffer (30% polyethylene glycol) during a 30 minute centrifugation step for precipitation. The supernatant was removed and the DNA/RNA pellet was washed with 11.98 mol/L ethanol, cooled at −20 °C. Following a final centrifugation (13 000 rpm, 4 °C, 4 mins), the ethanol was removed and evaporated at room temperature (20°C). Finally, the DNA/RNA was resuspended in 0.01 mol/L EB buffer from the QIAGEN kit. The aliquots intended for DNA amplicon sequencing were stored at −80 °C. The aliquots intended for RNA amplicon sequencing were further processed by first removing the DNA with the TURBO DNA-free Kit (Thermo Fisher Scientific) according to the manufacturer’s protocol. The RNA was then converted into cDNA with the RevertAid First Strand cDNA Synthesis Kit (Thermo Fisher Scientific). The extracts were stored at −80 °C, until being sent, together with DNA extracts, for amplification and sequencing of the V9 region of the 18S rDNA, with the 1380F/1510R primers, based on the Pjevac et al. (2021) protocol.

The bioinformatic processing of the raw sequence data was done using DADA2 (Callahan et al., 2016) to select the viable amplicon sequence variants (ASV). The processing was performed by the Joint Microbiome Facility (JMF) under the project ID JMF-2301-09. Sequentially, quality filtering was performed, as well as forward and reverse read merging, and further processing of the ASV abundance table by removing the ASVs that could not be classified at the supergroup level or those that were higher level eukaryotes. Sequence classification was based on the PR2 database (Vaulot, 2023).

### 2.4 Statistical analyses

All data was analyzed and visualized in R (R Core Team, 2024; Wickham, 2016). Sampling sites with sequence count *>*100 made up 46 DNA-analysed and 43 RNA-analysed samples. Species accumulation curves, diversity estimates and principal component analysis of the environmental parameters were performed with “vegan” functions under default settings (Oksanen et al., 2001). The community turnover and seasonal dissimilarities was calculated as Euclidean distances of central log-ratio transformed counts with “microViz” (Barnett et al., 2021). Distributions and spread for univariate variables were tested, and compared between groups according to the assumptions, with “rstatix” (Kassambara, 2023), while community composition was tested with “ microViz” wrappers (Barnett et al., 2021) for “adonis2” and “betadisper” functions. The relationship of community composition to significant environmental variables was identified (“capscale”) and tested (“anova.cca”) with “vegan”, after filtering out variables with a variance inflation factor (vif) *>* 10. Seasonality was tested with the combined test from the “seastests” package (Ollech and Webel, 2020), and differential analysis and effect size estimation was performed with microeco’s lefse function (Liu et al., 2021; Segata et al., 2011).

## 3 RESULTS

### 3.1 Sampling site characterization

Both the aquatic habitat type (surface water vs groundwater) and season shaped the environmental conditions (Fig. 2). The groundwater sampling site, termed D15, was characterized by elevated CO_2_ concentrations, DIC concentrations, sulfate and ortho-phosphate, in comparison to the surface water, ESW (t_CO2_ = 7.77***; W_DIC_ = 403**, W_SO4_ = 350**, and W_Ortho-P_ = 455***), Tab. 1. The microbial activity, measured as intracellular ATP, and prokaryotic cell densities were significantly lower in groundwater (W_ATP_=0***, W_TCC_=0***), as were the DOC and DO concentrations, pH levels, and ammonium concentration (W_DOC_ = 137***; t_DO_ = −8.7***; t_pH_ = -5.82**, t_NH4_ = −2.5***).

**Figure 2.**
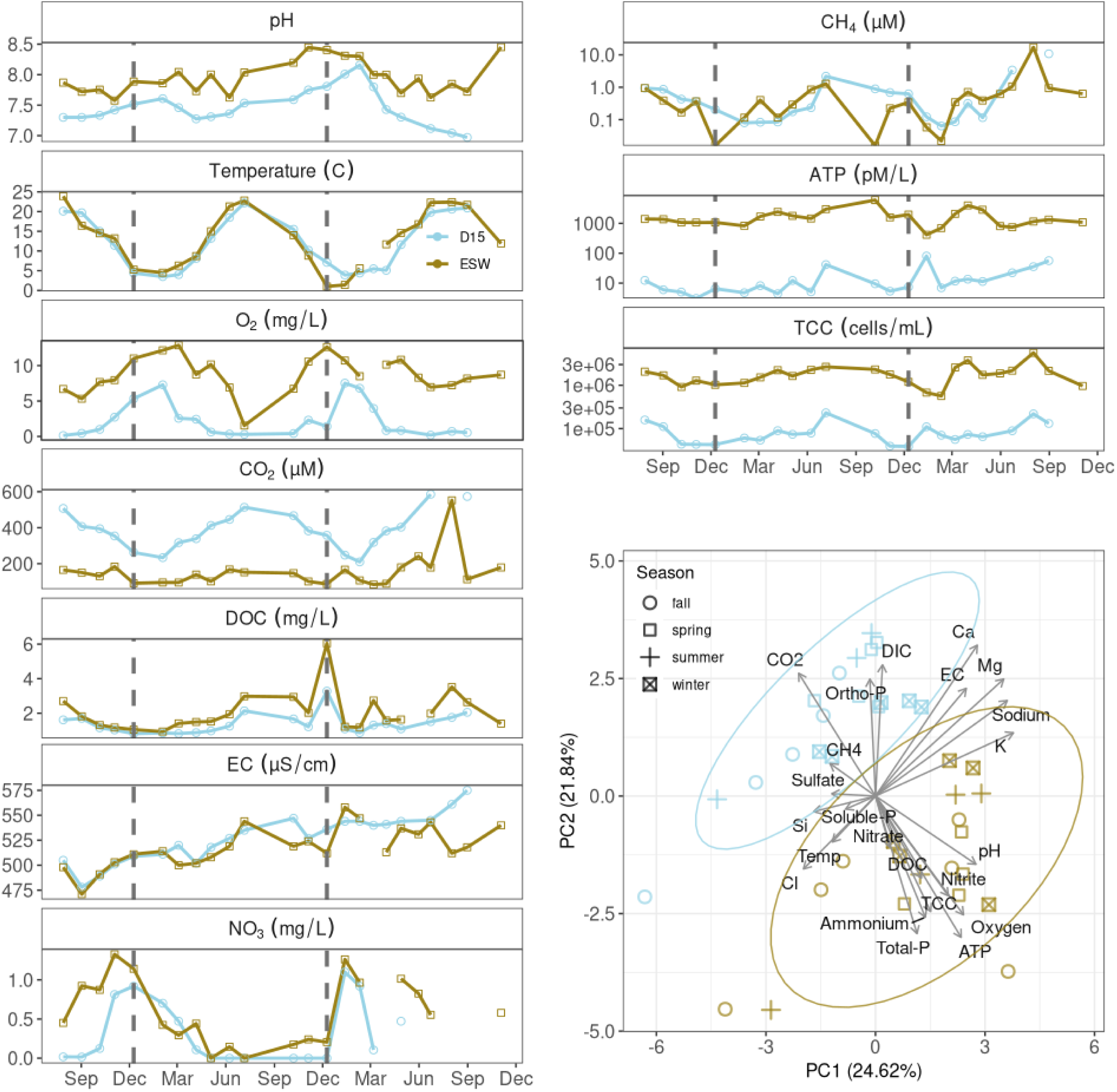
Time series of dynamic environmental variables for summer 2020 to winter 2022. Dashed lines represent the start of new year. Environmental characterization of the surface water and groundwater sampling sites via PCA. Color key: blue corresponding to the D15 groundwater sites, brown to the ESW surface water. ATP: internal ATP [pM/L], CO2: carbon dioxide [*µ*M], DIC [mg/L], DO: dissolved oxygen [mg/L], DOC: [mg/L], CH_4_: [*µ*M], Electric conductivity [uS/cm], TCC: [cells/mL], Sulfate: [mg/L], Ca: calcium [mg/L], Mg: magnesium [mg/L], Sodium [mg/L], K: potassium [mg/L], Ortho-P: orthoposphate [*µ*g/L], Si: silicate [mg/L], Temp: [°C], Soluble-P: soluble phosphate [*µ*g/L], Total-P: total phosphorous [*µ*g/L], Cl: chloride [mg/L], Ammonium: [*µ*g/L], Nitrate: [*µ*g/L]

**Table 1.** Values for environmental variables, means and standard deviations written by sampling site.

| Environmental variable | Site ID | Value (mean ± sd) |
| --- | --- | --- |
| pH | D15 | 7.47 ± 0.30 |
| | ESW | 7.96 ± 0.27 |
| Temperature [°C] | D15 | 12.03 ± 6.75 |
| | ESW | 13.22 ± 7.13 |
| Dissolved oxygen [mg/L] | D15 | 2.21 ± 2.43 |
| | ESW | 8.72 ± 2.59 |
| Carbon dioxide [μM] | D15 | 386.43 ± 104.75 |
| | ESW | 154.32 ± 94.13 |
| DOC [mg/L] | D15 | 1.39 ± 0.58 |
| | ESW | 2.05 ± 1.13 |
| DIC [mg/L] | D15 | 57.28 ± 4.98 |
| | ESW | 53.59 ± 4.65 |
| Electric conductivity [μS/cm] | D15 | 527.23 ± 23.80 |
| | ESW | 518.04 ± 20.45 |
| Silicate [mg/L] | D15 | 2.17 ± 0.41 |
| | ESW | 2.02 ± 0.57 |
| Nitrate [μg/L] | D15 | 0.32 ± 0.4 |
| | ESW | 0.59 ± 0.42 |
| Nitrite [μg/L] | D15 | 0.85 ± 2.17 |
| | ESW | 1.80 ± 0.91 |
| Ammonium [μg/L] | D15 | 10.60 ± 10.39 |
| | ESW | 19.60 ± 13.56 |
| Sulfate [mg/L] | D15 | 27.73 ± 4.73 |
| | ESW | 25.94 ± 1.94 |
| Orthophosphate [μg/L] | D15 | 2.27 ± 1.63 |
| | ESW | 0.53 ± 0.47 |
| Soluble phosphate [μg/L] | D15 | 3.16 ± 1.74 |
| | ESW | 2.71 ± 1.24 |
| Total phosphorous [μg/L] | D15 | 4.39 ± 3.28 |
| | ESW | 16.96 ± 20.29 |
| CH<sub>4</sub> [μM] | D15 | 7.04 ± 28.04 |
| | ESW | 1.14 ± 3.42 |
| Internal ATP [pM/L] | D15 | 16.67 ± 19.85 |
| | ESW | 1745.36 ± 1233.71 |
| TCC [cells/mL] | D15 | 8.91 $\times 10^4 \pm 5.38 \times 10^4$ |
| | ESW | 1.90 $\times 10^6 \pm 1.07 \times 10^6$ |

Seasonal dynamics in CO_2_, DOC, DO, pH, and temperature were apparent at both sampling sites, but not significant (Ollech and Webel seasonality test showed all variables to have p*>*0.05). Groundwater temperatures showed a short time-lag following the surface water dynamics, but were comparable in amplitude ranging from (22.5°C) in summer, to 1 °C in winter. The gasses (CO_2_, DO and CH_4_) followed a general yearly trend, but with considerable differences in amplitudes. In ESW the DO was mostly above 6 mgL^-1^ (8.72 ± 2.59) with the exception (1.54 mgL^-1^) of July 2021, while groundwater from D15 was predominantly hypoxic (2.21 ± 2.43). Groundwater samples exhibited more dynamic CO_2_ concentrations than surface water samples, with peaks in summers, when concentrations of dissolved O_2_ decreased to hypoxic and anoxic conditions. However, surface water was more variable than groundwater when it came to methane levels, and showed sudden declines in winters. Dissolved organic matter and pH varied throughout the year, but without a straightforward summer-winter contrast observed for other environmental variables. Still, similar temporal trend could be noted for both sampling sites.

Electrical conductivity of the two sites differed little in winter 2020, with groundwater conductivity only stabilising as higher than surface water towards the end of our study period, in 2022. Values of *δ*^18^O and *δ*^2^H in water showed comparable isotope trends (*ρ*_18O_ =0.94, *ρ*_D_ =0.93) for both sites, but with a higher deuterium excess permil in late summer, fall and winter 2020 (10.2 ± 0.81) than in spring and summer 2021 (9.26 ± 0.43). The groundwater and surface water *δ*^18^O values were in close proximity (Fig. S1, left), and showed surface water strongly influencing the shallow aquifer. In other words, the porportion of surface water infiltrated in the shallow aquifer was high. This was also supported by the water level differences for almost all time points (Fig. S1, right). Isotope measurements for year 2022 were not available and could not be used for a comparison to the changes in electric conductivity behaviour.

The prokaryotic cell densitites (TCC) and microbial activity (ATP) exhibited drastically different values and ranges between the two sites. Both measures were lower by two orders of magnitude in groundwater than in surface water (Tab. 1). Overall, the similarity in trends was discernable, despite the differences in absolute values. The summer peaks in prokaryotic activity and abundance were, however, clearer in groundwater.

### 3.2 Diversity and richness of protist communities

Species accumulation curves indicated that protist communities in neither surface water nor groundwater were fully captured with the current sampling effort, since neither DNA- nor RNA-detected species reached a plateau (Fig. 3, top). However, in comparison to surface water, groundwater showed a steeper slope for both barcoding methods, and a comparison of DNA- and RNA-based richness showed that the DNA-based method detected a slightly greater number of taxa. In DNA- and RNA-derived communities, 3,368 and 3,100 taxa were present, respectively. Not only was the richness of detected ASVs higher in surface water, but the sequence counts also showed pronounced differences by site (DNA: median_(D15)_=8,187, median_(ESW)_=10,148; RNA: median_(D15)_= 6,173, median_(ESW)_=13,982).

**Figure 3.**
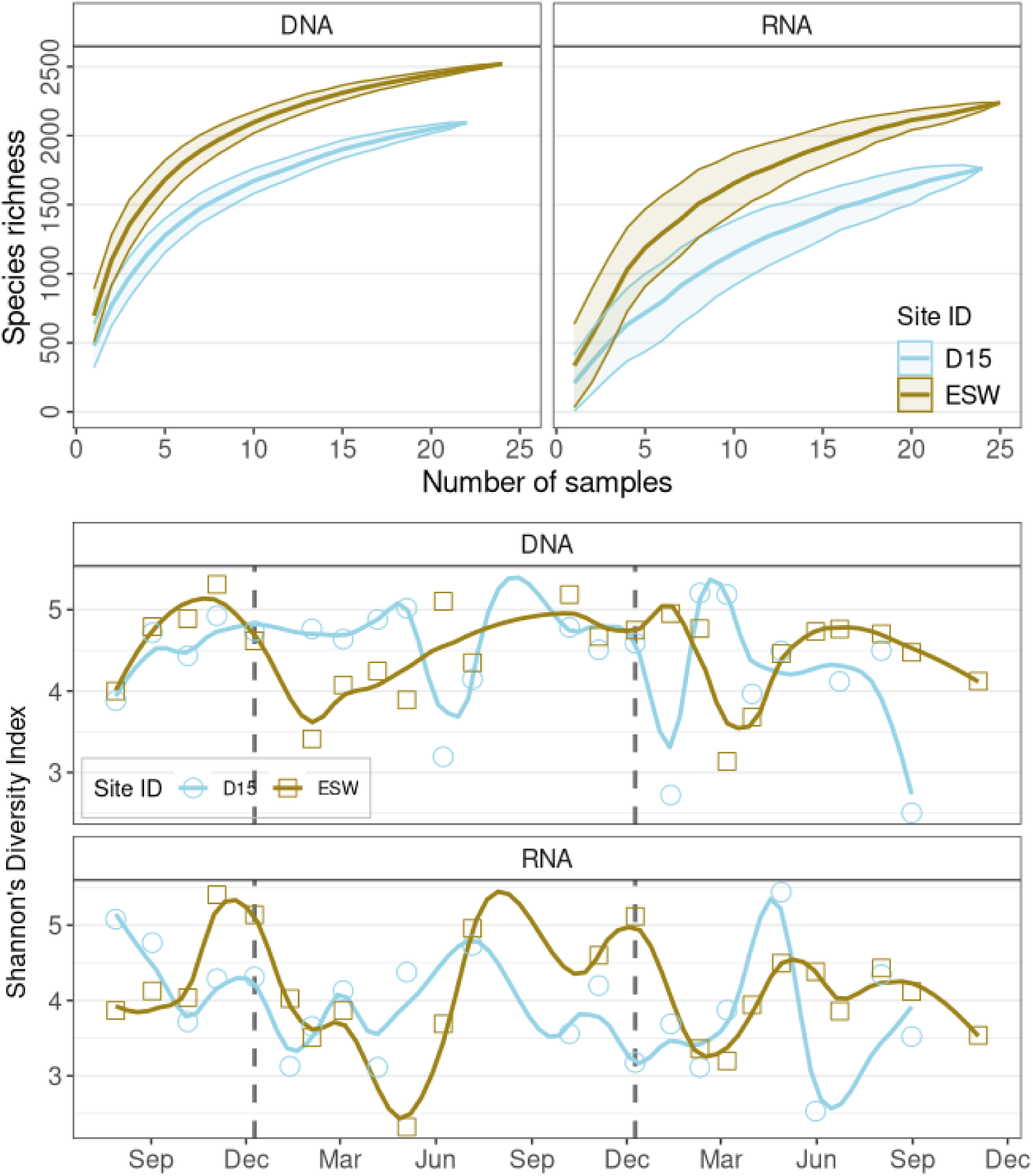
**Top.** Species-accumulation curve for the samples, according to the sampling site and method. **Bottom.** Protist Shannon’s diversity index with a loess fit, according to sampling date and method.

The higher richness in surface water did not translate to a higher Shannon diversity (Fig. 3, bottom). There was no difference between the surface water and groundwater in Shannon diversity, but seasonality, especially for the DNA-derived community, when accounting for the site specificity, was significant (F_(7, 38)_=2.59, p=0.028, adj. R^2^ = 0.198). Season as the only factor, or site alone, could not be identified as significant drivers of diversity. The lowest protists diversity in surface water can be seen in spring, when peak diversity was observed in groundwater. The most similar values between the sites could be seen in the winter. The active community exhibited a more dynamic behaviour throughout the year, with sites having the highest similarity in springs.

The season as a category explained the diversity behaviour better than any combination of the environmental variables recorded, including those that behaved seasonally, but significant trends showed surface water diversity following the behaviour of ammonium (*ρ*=-0.68,) and nitrate (*ρ*=0.48), and groundwater diversity corresponded to changes in temperatures (*ρ*=0.56), dissolved oxygen (*ρ*=-0.51), pH (*ρ*=-0.46), CO2 (*ρ*=0.58), CH4 (*ρ*=0.58), and microbial activity (ATP) (*ρ*=-0.23).

### 3.3 Seasonal dynamics of communities

Groundwater active communities experienced significantly higher turnover than its DNA counterpart (F_(1,41)_=6.34, p*<*0.02), but there was no difference in turnover between the sampling sites. Community composition was significantly different for groundwater and surface water, for both the DNA (F_(1, 44)_=14.34, p=0.001, adj.R^2^=0.25) and the RNA extracts (F_(1, 41)_=6.22, p=0.001, adj.R^2^=0.13). The effect of seasonality was strong for both sites and both DNA and RNA based communities, showing composition to follow a yearly cycle from summer- to winter-specific assemblages, including the fall/spring transitions (Fig. 4). Season alone explained 5.3% of variance in the DNA community, and together with habitat type and the interactions between them, reached 34.5% (F_(7, 35)_=4.16, p*<*0.001). The return to the summer composition was even clearer when following community dissimilarity through time, where we saw compositions of summers 2021 and 2022 more similar to the composition of the starting point in August 2020, and strongly differing from winter compositions (Fig. 5). The dissimilarities between the sites remained stable throughout the whole period of study, and were higher than any within-site dissimilarity. Total community could be tied to seasonally-dependant environmental variables such as temperature, microbial activity and CO_2_ levels (F_(3, 39)_=5.9, p*<*0.001, adj.R^2^_DNA_ = 0.259), but these variables explained less composition variability than season and site constraints. Active communtities were also tied to DOC, NO_3_, and CH_4_ (F_(3, 36)_=2.37, p*<*0.001, adj.R^2^_RNA_ = 0.10), compared to 15% variance explained by the season and site (F_(7, 32)_=1.99, p*<*0.001).

**Figure 4.**
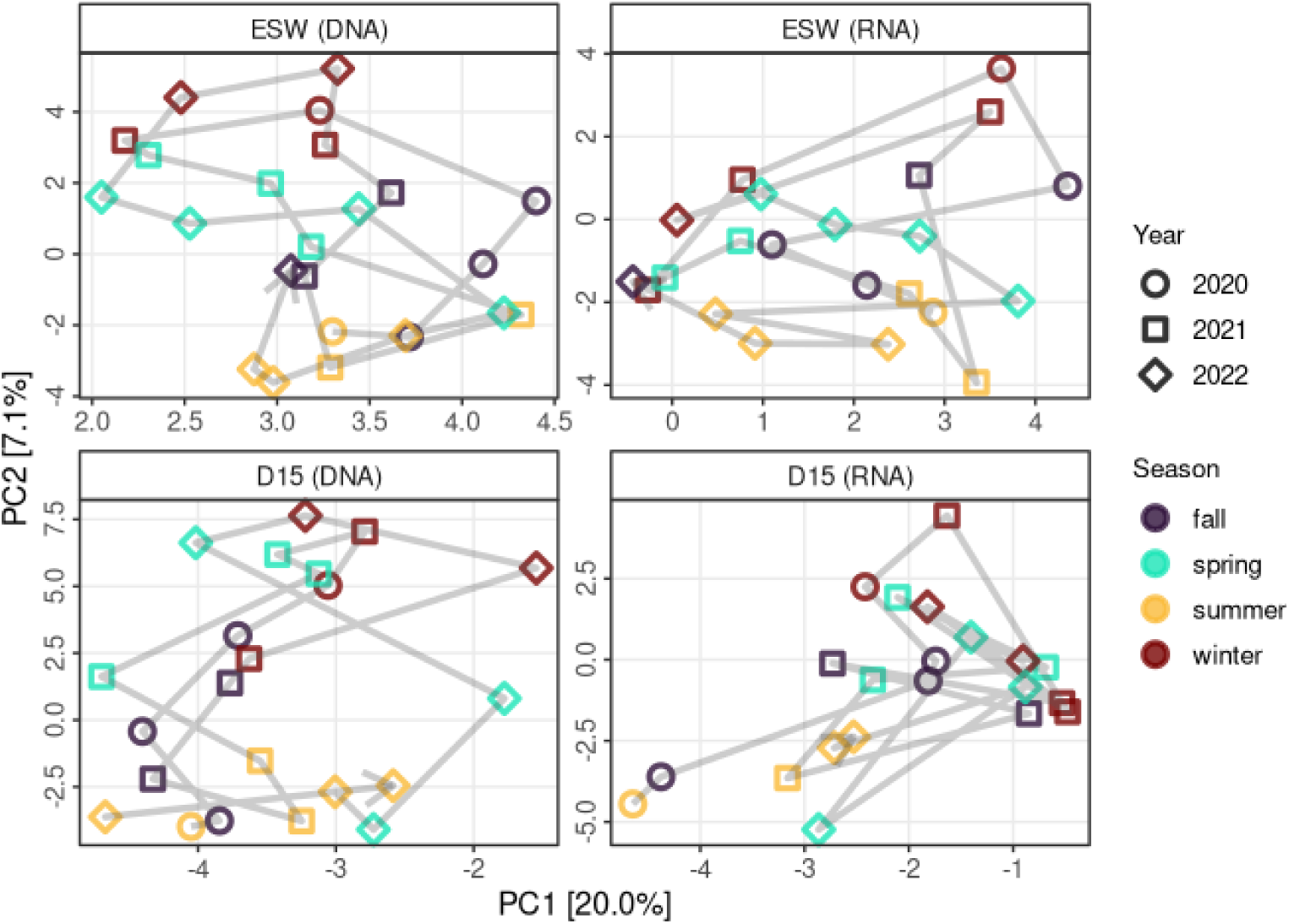
PCA of the protist composition for each sampling site and method used, according to the season and year. The sampling times are represented as a sequential line between the points.

**Figure 5.**
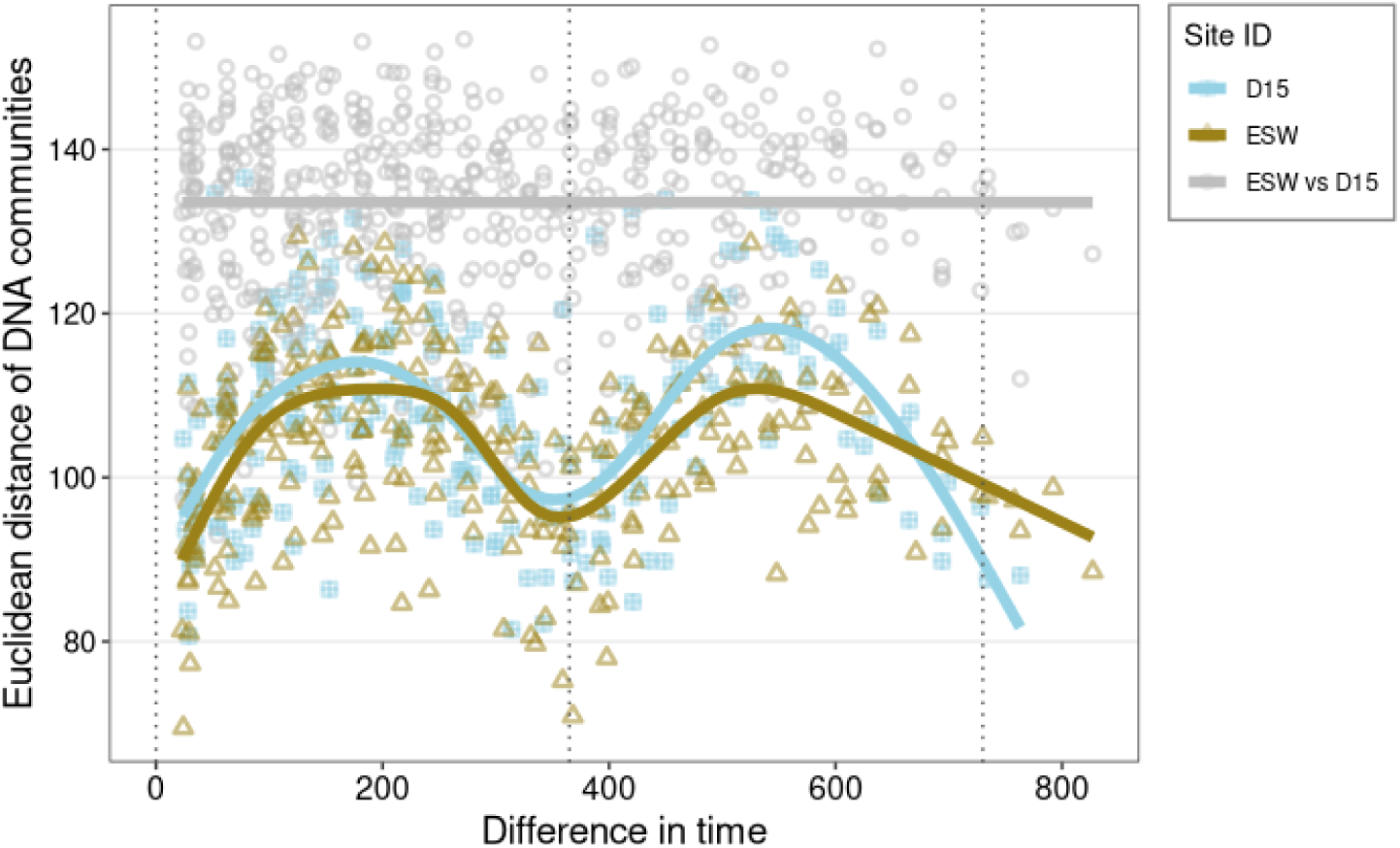
Community pairwise dissimilarity of clr-transformed abundance counts according to Euclidean pairwise time distance in days, colored by sampling site, with the loess fit in corresponding color.

### 3.4 Community composition

The distribution of ASVs showed that taxa shared between the two sites were not much less numerous than the number of ASVs specific to each habitat (Fig. 6). However, abundance-wise the shared ASVs were the more dominant part of the communities. For DNA-derived communities, 68.6% of the total relative abundance corresponded to ASVs that were shared between surface water and groundwater, and similar pattern was observed for RNA-derived communities, where 62.4% of the relative abundance was attributed to shared ASVs.

**Figure 6.**
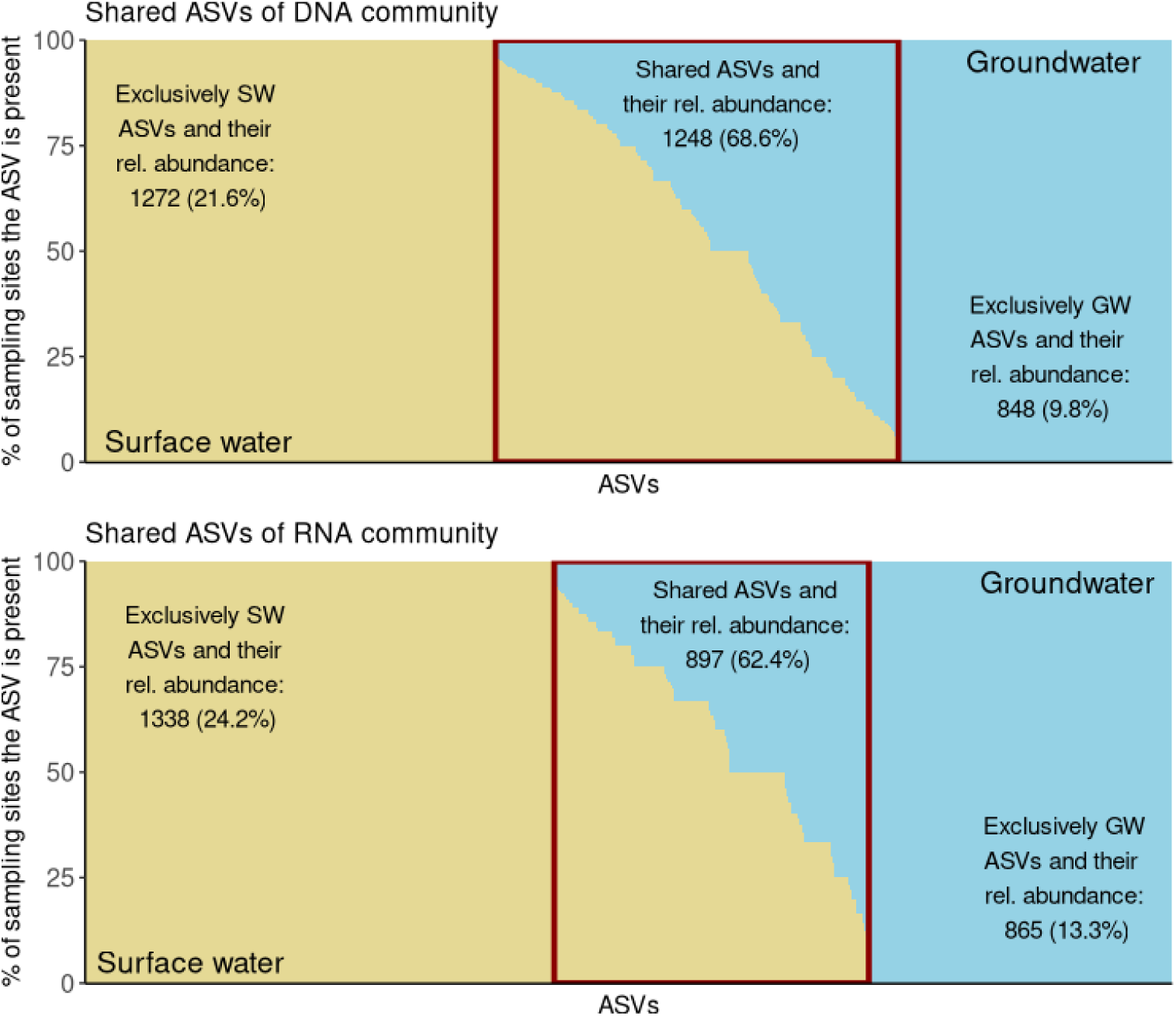
ASV distribution in groundwater and surface water, with red rectangle denoting the shared ASVs. Numbers of taxa and their relative abundances in the habitat they were present in are written for each fraction.

Ochrophytes and ciliates were the most abundant protistan divisions at both sampling sites. Higher relative abundances of ochrophytes in total and active communtities were observed in surface water (ESW_(DNA)_=39.96% ± 10, ESW_(RNA)_=50.52% ± 23), than in the groundwater (D15_(DNA)_=23.36% ± 18, D15_(RNA)_=24.15% ± 27), whereas ciliates were more inclined to groundwater, making a big portion of active community (38.84% ± 23.2). The median ratios of ciliates to ochrophytes were higher in groundwater for both methods, but the difference between two sites was more drastic for active communities (D15_DNA_=0.75, ESW_DNA_=0.43; D15_RNA_=2.7, ESW_RNA_=0.36).

*Spirotrichea* were observed as both the most active and most abundant ciliates in surface water, whereas *Oligohymenophorea* (the classified ones mainly belonging to genus *Tetrahymena*) made up a significant portion of the groundwater ciliates (Fig. 7). *Ciliophora* in groundwater were also more likely to be unclassified, than ciliates found in surface water. *Chrysophyceae* made up a big portion of total communities’ ochrophytes at both sites, but the most active *Ochrophyta* were *Bacillariophyta*. Both *Chrysophyceae* and *Bacillariophyta*, together with *Dictyochophycaeae* and *Synurophyceae*, showed stronger inclination towards surface water, while *Eustigmatophyceae* could be designated as groundwater ochrophytes (Fig. 8). The *Bacillariophyta* in question were mainly *Navicula* and *Staurosira*. Flagellates, such as cryptophytes *Prymnesiophyceae* and *Cryptophyceae*, as well as chlorophytes, choanoflagellates and *Katablepharidaceae* were associated with surface water. In terms of presence, *Cercozoa* were found in both surface water and groundwater. However, surface water was characterized by the cercozoan *Filosa Thecofilosea*, while groundwater had *Filosa Sarcomonadea* as the most pronounced *Cercozoa* class. Although *Filosa Thecofilosea* were present only in low abundance in groundwater, they were composed predominantly of active taxa. Taxa which were significantly more inclined to groundwater in terms of both presence and activity, other than the mentioned ciliates and *Filosa-Sarcomonadea*, were *Euglenozoa* and *Apusomonadidae* Group 1.

**Figure 7.**
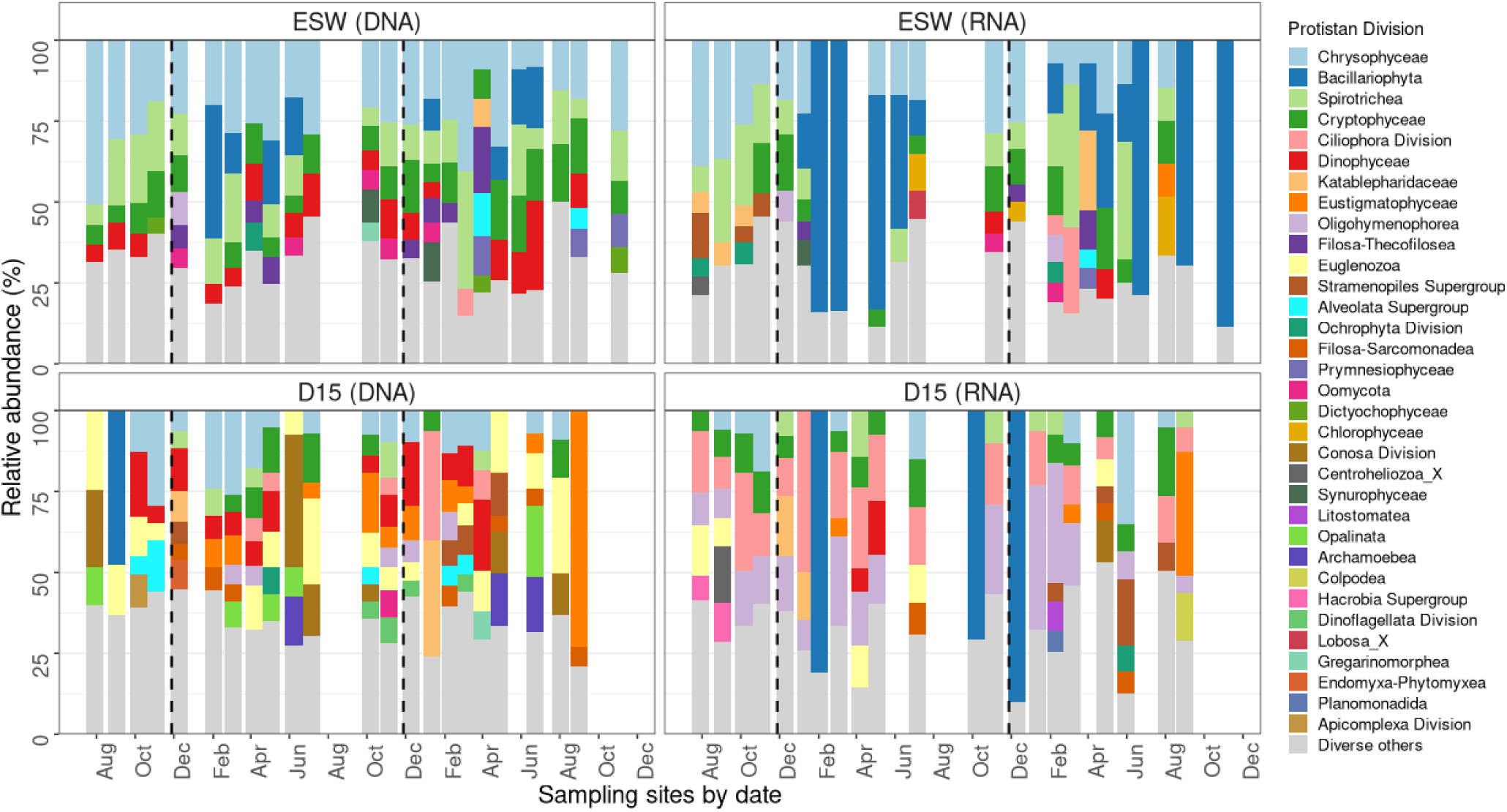
Main protist divisions and their relative abundance in surface water (ESW) and groundwater (D15), according to the barcoding method. Taxa with *<*5% of relative abundance is classified under “Diverse others”.

**Figure 8.**
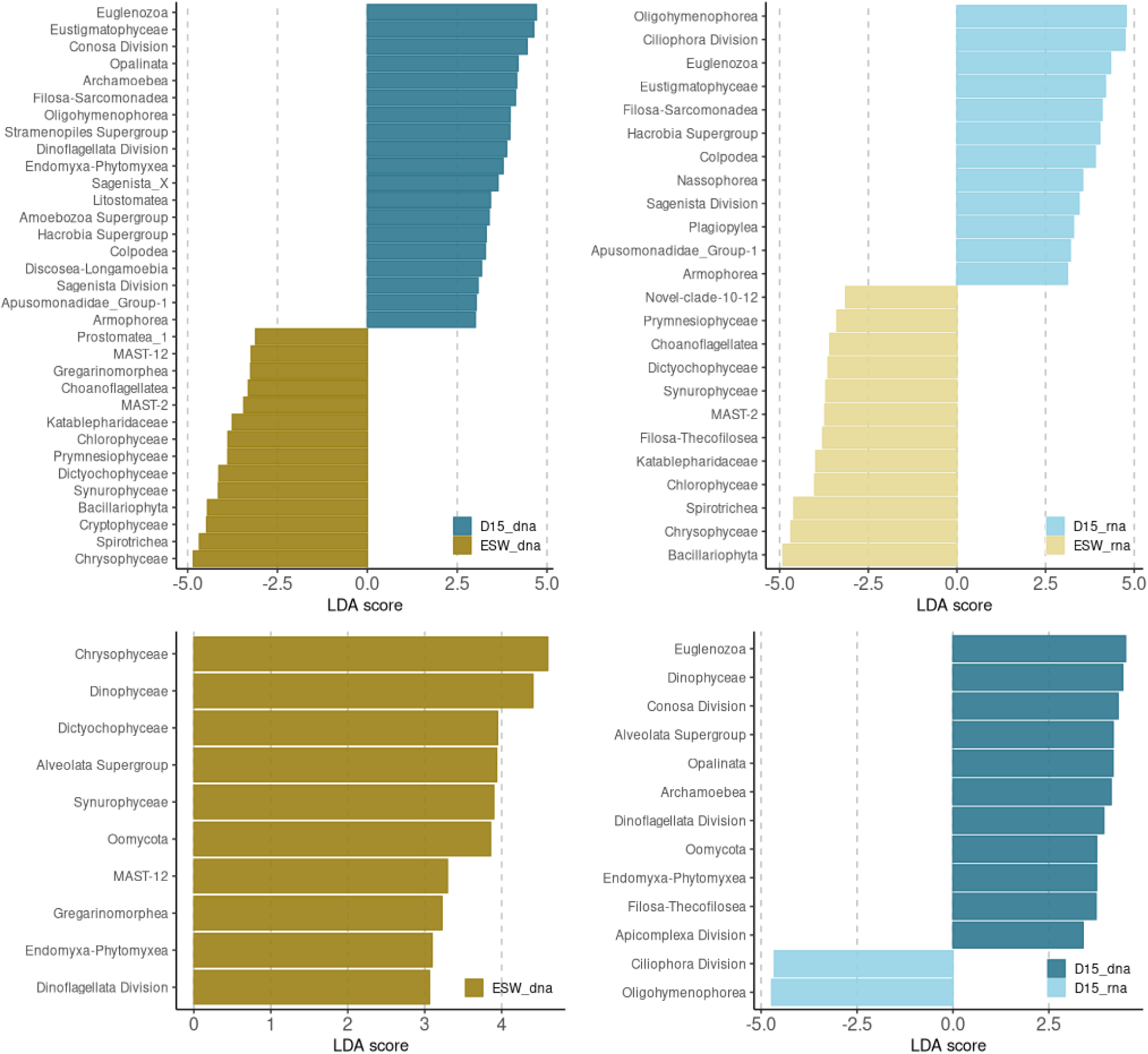
Differential abundance analysis by Linear discriminant analysis Effect Size (LEfSe) of the total and active communities. The bar plots show significant taxa with the effect size *>*3.

Five sampling time points characterized by sequence counts drastically different from the general distribution (30-40 sequence counts) were considered separately and showed differing composition between themselves and compared to the remaining samples (Fig. S2). Taxa which dominated these few samples were usually noted as rare taxa only sporadically present in the rest of the samples. Most notably, groundwater composition of March 2022 was dominated by *Gregarinomorphea*, absent from most other sampling points.

### 3.5 Seasonal trends of individual taxa

Taxa associations to specific yearly periods were present for both winter and summer (Fig. 9). *Archamoebea* and unclassified *Conosa* showed inclination to summer peaks in groundater, while their abundances remained stable in surface water. *Telonemia* abundances peaked in early summer in surface waters, whereas unlassified *Apicomplexa* and protists of Novel clade 10-12 reached their surface water peaks in falls. Winters and springs were optimal periods for *Filosa Thecofilosea* in both groundwater and surface water, but *Katablepharidaceae* remained stable in surface water throughout the year, while still exhibiting winter peaks in total and active communities of groundwater. Other groundwater-specific seasonal trends were observed for *Pirsonia* clade, as well as for *Apusomonadidae* Group 1 and *Bolidophycae*, which both showed a lag in active community in comparison to total community. CONTH 4 showed groundwater peaks which followed the surface water winter peaks, but with limited effect on active groundwater commuinities. Lastly, *Oligomenophorea* showed clear winter peaks in surface water, but experienced more drastic variablity in active groundwater communities.

**Figure 9.**
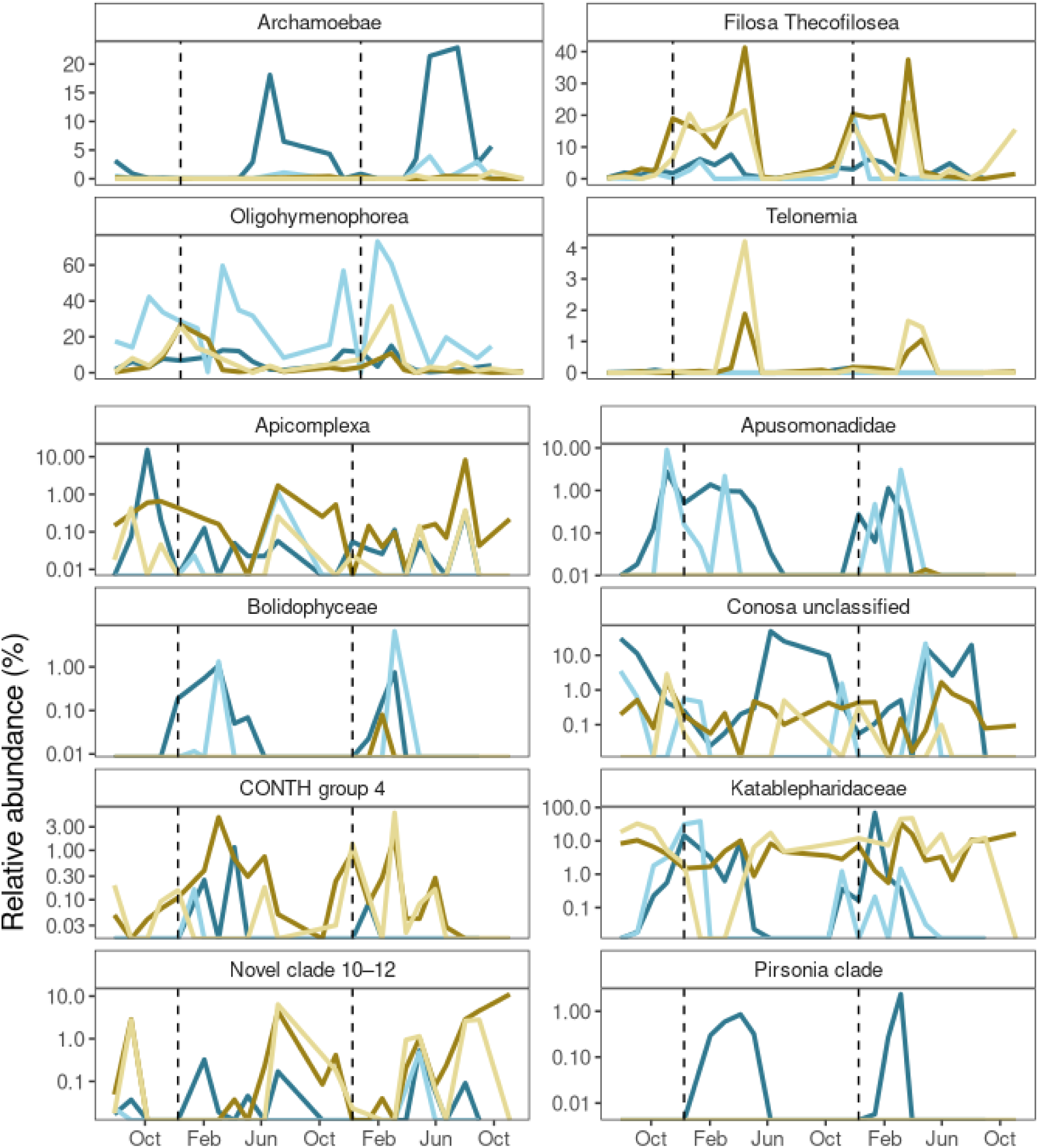
Seasonal dynamics of protistan taxa in groundwater and surface water over a two-year period. Relative abundances (%) protistan classes which showed significant differences between seasons shown across sampling dates. Line colors correspond to different sampling site and extraction method. Color key: blue corresponds to the D15 groundwater sites, brown to the ESW surface water, lighter shades represent RNA extracts, and darker the DNA extracts. Dashed vertical lines indicate year start.

## 4 DISCUSSION

Scarcity of studies on groundwater protists, as well as a lack of research conducted by applying state of the art molecular techniques leave a large gap in our knowledge of subsurface eukaryote communities and its dynamics. A first overview of groundwater protistan communities in surface water and groundwater of the Danube Floodplain National park, Austria, is provided by sequencing the V9 region of the 18S rDNA marker gene. Motivation for this study was to look into the transitional change between a surface water and groundwater habitat and determining the hypothesized site-specificity, i.e. lack of light and limitations of resources impacting composition, functionality and diversity of protist communities. To our knowledge, this is the first study that focuses in detail on the protist community composition and turnover of a groundwater and nearby surface water habitat for two hydrological seasons.

### 4.1 Site specificity and environmental filtering

The shallow groundwater monitoring site in the Danube wetlands was chosen as a suitable shallow subsurface site for exploring seasonality and surface water influence on protistan and prokaryotic communities. Trends in water levels and stable water isotope signatures, in the period with data available, indicate groundwater recharge by surface water most year round. Dependent to the difference in hydraulic heads, the water traveling time from the backwater to the shallow groundwater site sampled is assumed to take 2-4 weeks for the 25 m distance (Dreher et al., 2006). A continuous hydrological connectivity was supported by predictable patterns in temperatures, dissolved oxygen and carbon dioxide throughout both years. The summer periods were characterised by higher temperatures both in surface water and groundwater, which promoted microbial activity and growth, resulting in increased respiration rates, declining oxygen and releasing CO_2_, in a manner previously observed for this monitoring site (Dreher et al., 2006). The depleted nitrate with methane spikes indicated a period of reduced conditions in the surface water sediments and in the aquifer in summers, which started changing towards a more oxygenated habitat only in late winters.

The contrasting trends of continuous changes in most redox species versus the differences in electrical conductivity and DOC for the two years of the study, were considered for the protistan community analysis. With the differences in conductivity and the spikes in organic matter between the two years, it was likely that sudden recharge events and the follow-up disturbance could be a driving force of protistan community dynamics. In this case, the year of sampling would be a good predicting factor. This was not observed, since the differences between groundwater and surface water sites remained stable throughout all seasons, for both years. Both total and active communities were site-specific in composition, indicating that even the fraction of the community that persists long enough to be detected as DNA extracts, regardless of its ability to be active in groundwater, was distinct from the surface community. Whether this could be due to selective attenuation due to size, shape or other travel-related proccesses, needs to be assessed in future studies, but for the time being, it seems not to be due to periods of increased recharge. Additionally, most season-specific taxa also showed similar behaviour for both years, regardless of differences in recharge probability.

While difficult to decouple seasonality in shallow groundwater, in terms of temperature and temperature-dependent processes, from the effects of seasonal recharge patterns, the lack of a detectable relationship with electrical conductivity on protistan communities, which can, in some cases, act as a proxy for surface water infiltration in some cases, combined with the influences of CO_2_, CH_4_, and NO_3_, indicate a close relationship with seasonal changes in biogeochemical processes, which could be only indirectly impacted by recharge. Ultimately, the temporal community gradient was stable, reaching the starting point after each yearly cycle, implying that despite the connectivity between the two sites, be it continuous or sporadic, it is the specificity of shallow groundwater habitat that was the likeliest abiotic driver of protistan communities during the study period. The higher similarity between the two sites’ communities in winters, which we hypothesised would be the case, was not proven valid. The theorized less pronounced seasonal dynamics in groundwater communities in comparison to surface water protists was disputed, as well. The pronounced seasonality observed at this groundwater site could be attributed to its shallow depth, which translates into a close link to temperature fluctuations occurring on the surface and in the surrounding soil, even in the absence of direct groundwater recharge. The upper soil layers would be a source of labile, bioavailable dissolved organic matter, in comparison to the refractory DOM in deeper soil layers (Kaiser and Kalbitz, 2012). The temperature increases, in combination with bioavailable DOM input, could stimulate microbial growth and activity, and cause shifts in biogeochemical proccesses rates during summer months (Brielmann et al., 2009). While presence of phtotrophic protist taxa in groundwater supports the idea of a continuous connectivity to surface water, it is likely that seasonality of protistan communities in groundwater is strongly exacerbated by the recharge-independent seasonality.

### 4.2 Groundwater-specific protist taxa

The taxonomic overlap between groundwater and surface water suggested that in both active and total communities, it is taxa shared between the two sites which prosper, despite the assumption of groundwater habitats selecting for a highly adapted community. This is likely due to the short distance between the water bodies and due to selective pressures brought on by the continuous hydrological connection between the surface water and groundwater. The site-specific community composition was shown primarily in the differing ratio of ciliates and ochrophytes, with most groundwater-specific taxa having cillia as locomotion system (such as *Oligomenophorea*, *Colpodea*, *Nassophorea* and *Armophorea*). Although ciliate abundance can be questioned due to molecular biases mentioned previuosly (Medinger et al., 2010), our study implemented the same sampling, extraction and sequencing protocols for groundwater and the near-by surface water, making it comparable within the study, and giving confidence to the site-specific ciliate abundances and activity. Additionally, *Cilliophora*, while more abundant in groundwater, were not the dominating taxa there - protists using flagellas for mobility dominated the groundwater site, with amoebae and ciliates making a smaller portion of all taxa. This fits the previous reports of groundwater protistan communities (Risse-Buhl et al., 2013; Loquay et al., 2009), and the lack of ciliates could, as per interpretation of Novarino et al. (1997), partly be a result of aquifer geology and pore size, as well as trophic interactions. With the sand aquifer being dominated by flagellates Novarino et al. (1997), and the karstic groundwater system predominately inhabited by ciliates (Herrmann et al., 2020), it is possible that the more even ratio of ciliates and flagellates in the shallow aquifer of our study can be a result of the mixed coarse gravel, sand and silty-clay layers (Danielopol, 1989). This relationship between aquifer characteristics and protist community composition is yet to be further explored and confirmed in further studies. Still, it remains questionable whether groundwater habitats provide enough trophic resources to sustain larger populations of ciliates and amoebae over time, regardless of aquifer characteristics. The abundant bacterial prey, which these protists require for survival, have a significantly lower biomass in groundwater compared to surface water (Griebler and Lueders, 2009), and future studies should aim at quantifying all members of the microbial food web in order to override the doubt of molecular bias and to better understand the conditions under which such taxa persist and remain active in subsurface ecosystems.

It is important to note that the study focused solely on the planktonic portion of the protistan community, and did not include barcoding of the attached or biofilm-associated taxa. This limitation could also have contributed to the excessive presence of ciliates in the free-living community of groundwater. In karstic aquifers, ciliates are indicated to dominate under anoxic conditions, while cercozoans only become competitive once oxygen levels increase (Herrmann et al., 2020). A similar pattern was observed at our study site, where cercozoan *Filosa thecofilosea* was most abundant in winters and early springs, corresponding with *Oligomenophorea* abundances, and sufficient oxygen levels. Given the high abundance and activity, as well as the vast and well-established bioindicator potential of ciliate species (Foissner, 2004; Stoeck et al., 2018; Kulaš et al., 2021), and the site’s proximity to urban influences, long term monitoring should be considered in the future. In particular, the detection of *Tetrahymena*, which are associated with high saprobic conditions (Foissner, 2016), should be examined in the future.

Other groundwater-specific taxa included mainly those previously reported in other aquifers or soils, like *Apusomonadidae*, and euglenids, which survived even in aquifers contaminated with elevated levels of nitrate and chloride (Loquay et al., 2009), and soil-abundant cercozoan classes (Oliverio et al., 2020). The hypothesised scarcity of phototrophic taxa in groundwater was not immediately evident, due to the high fraction of ochrophytes, a predominantely phototrophic group in the RNA derived communities. However, the *Chrysophyceae*, *Eustigmatophyceae* and *Bacillariophyta*, have all shown mixotrophic potential under suboptimal conditions, as long as DOC levels were sufficiently high (Tuchman et al., 2006; Villanova et al., 2022; Lie et al., 2018). Overall, the dominant trophic guild in groundwater was heterotrophy, but with indications of a considerable amount of mixotrophic activity, and in summers, parasitic *Apicomplexa* showed peaks in activity. This supports the hypothesized general heterotrophic dominance in groundwater, while sporadic mixotrophic activity could reflect the previously mentioned transitional character of the site, caused by continuous connectivity to surface water.

Unclassified members of the *Conosa* group showed a notable increase in relative abundance in the DNA-derived communities in summers, although their activity, i.e. higher abundance in the RNA pool, was observed mainly in oxic periods, as would be consistent with previous reports for some members of the group (Cavalier-Smith, 1998). While tempting to designate differences in activity and presence as a result of dispersal from the surface, it could also be a consequence of pronounced cyst formation, a common strategy among protists under suboptimal conditions. Still, the lag in activity compared to when a DNA spike occurred, in winter-specific taxa *Bolidophyceae* and *Apusomonadidae*, could indicate that these protists are introduced into the groundwater from an external source, since both taxa appeared in total communities in early winter, but their activity was only observed a month later. Lastly, we found indications of taxa which showed divergent seasonal behaviors between the two sites, namely the seasonality of Novel clade 10 and 12. Their abundance peaked in summer in surface water, in a manner incompatible with groundwater. The current scarce knowledge on the ecology of this almost exclusively freshwater planktonic clade does not yet offer conclusive insights.

### 4.3 Diversity

Protist diversity of different realms spans a wide range of Shannon diversity values, which, combined with site-specific environmental differences, could provide grounds for hypothesising that diversity patterns between freshwater surface and groundwater habitats would differ. However, the unique challenges of subsurface conditions were just as likely to host a vastly diverse community, just being lower in total abundance and biomass. Protistan richness in shallow groundwater was indeed slightly lower than in surface water, but differences in Shannon diversity were not significant. The higher surface water richness may reflect greater external inputs of organisms, but could be a result of sequencing bias, since higher protist biomass in surface water could be translated to an increased sequencing depth (Schenk et al., 2019). Although cell quantification for eukaryotes was not within the scope of this paper, this would ideally be addressed in future research at these sites, focusing on dynamics between eukaryotes and prokaryotes, and relationships between the biomass of these two groups, as was done in Karwautz et al. (2022). For this study, biodiversity indices which consider the evenness might, therefore, serve as a more robust diversity estimate of amplicon data, than richness alone.

One of the few studies that dealt with protist diversity of transition zones between a high energy system and nutrient-poor habitat was conducted in a cave, at the point of inflow and at the point of stable no-light oligotrophic conditions. It reported a higher Shannon diversity at the surface water inflow point, where photosyntethic protists and organic matter were present (Gogoleva et al., 2024). As continuous hydrological connection cannot be excluded, a similar interpretation might be fitting for our study site, as a possible equivalent to the cave system, being a transitional zone with sustained environmental variability and colonisation opportunity. This could be the reason why groundwater from our sampling site did not deviate strongly from surface water communities in terms of diversity. In the future, we need to include in addition a deeper and more distant groundwater site assumed to host distinctly lower-diversity assemblages due to more stable and oligotrophic conditions.

The seasonal dynamics in the DNA-derived community diversity showed converging values of the two sites in winter, especially in December 2020, and further supported the idea of increased connectivity during this period. However, while diversity calculated based on DNA extracts converged in winters, active community diversity showed the opposite trend by greater winter divergences. Considering a lack of straightforward delay of groundwater diversity in active communities, and little comparability between the trends of two sites, decoupled diversity dynamics of the two sites was more likely than the connection between them. The differences in diversity drivers of active protistan communities supported this. While our two-year period was sufficient to capture community composition stability, longer-term monitoring might be necessary to elucidate the diversity dynamics, and its dependence on food web interactions.

## 5 CONCLUSION

Our study of protistan communities in the Danube Floodplain National Park, Vienna, Austria provides a detailed characterization of community composition, diversity, and feeding modes across a surface water to groundwater gradient, using molecular barcoding of total DNA-derived and active RNA-derived protistan communities. Monthly sampling over two hydrological years demonstrated a distinct and robust compositions and turnover of surface water and groundwater protistan communities. We showed that in terms of diversity active protistan communities in groundwater exhibited seasonal dynamics similar to surface water communities. Functionally, the groundwater community was dominated by heterotrophic and mixotrophic protists, and exhibited the presence and activity of ciliates. The groundwater protist diversity was connected to prokaryotic activity and redox conditions, and called for future screening of food web interactions. The study provides one of the first steps towards the molecular analysis of protist communities in groundwater. Long-term monitoring and inclusion of attached communities will be critical for refining our understanding of protist ecology in groundwater.

