## Supplementary Information (Fig S1-S4) for "Dynamic changes of protist community composition along a surface water-groundwater transect in the Danube wetland Lobau, Vienna, Austria"

### Supplementary Material

#### 1 SUPPLEMENTARY FIGURES

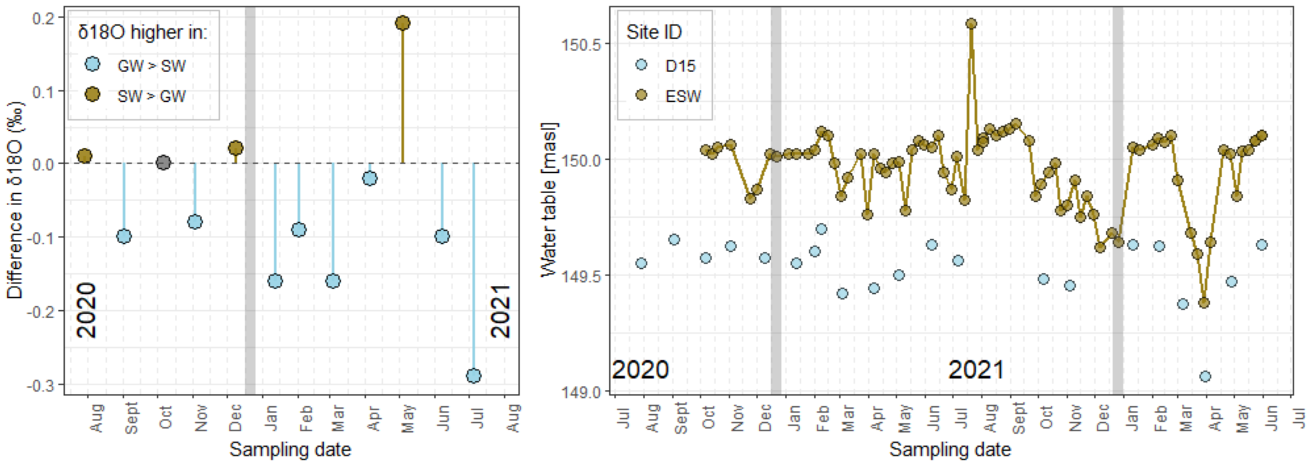


**Figure S1. Left.** Differences in $\delta^{18}$O values between surface water ESW and groundwater site D15, for period summer 2020-summer 2021. **Right.** Water levels of surface water and a nearby groundwater monitoring well in meters above the Adriatic, for period summer 2020 to summer 2022. Start of a new year is marked with a gray horizontal line.

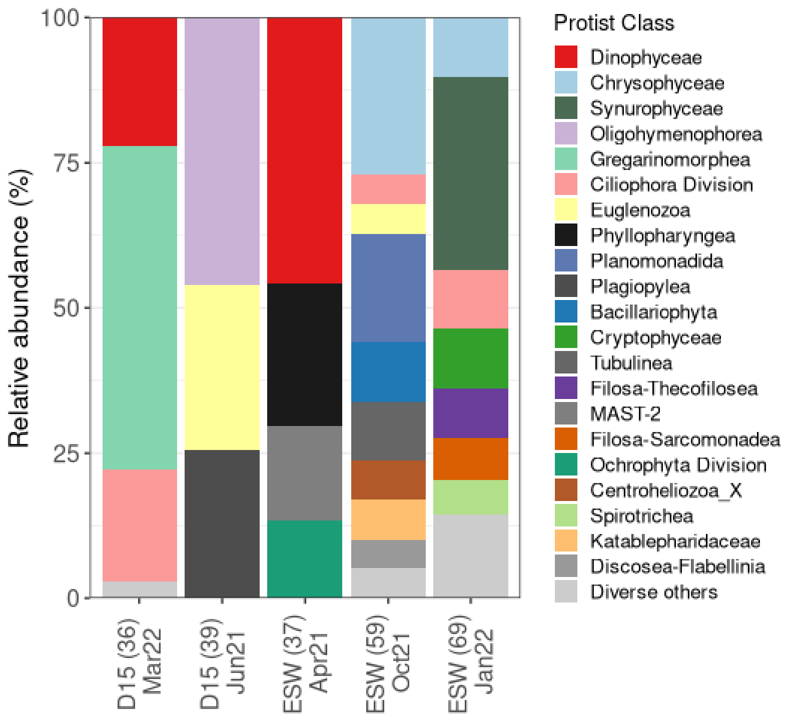


**Figure S2.** Relative abundances of classes identified in sampling points with <100 sequence counts. Sampling site and date of sampling together with read counts in paretheses are written on x-axis.

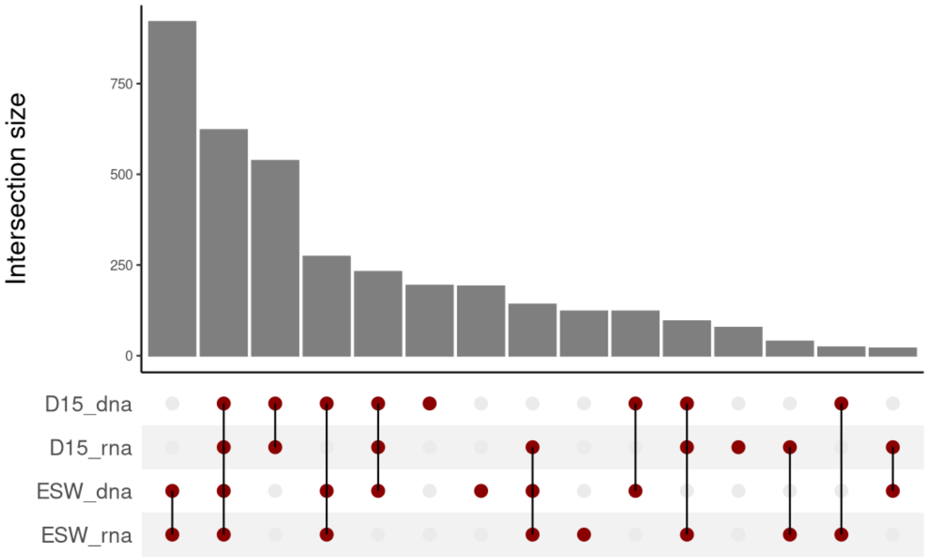


**Figure S3.** Shared protistan ASVs across the two sampling sites and two molecular methods. The bar plot on the top shows the size of ASV intersection across group combinations, the bottom indicates the groups involved in each intersection (dark red dots connected by lines).

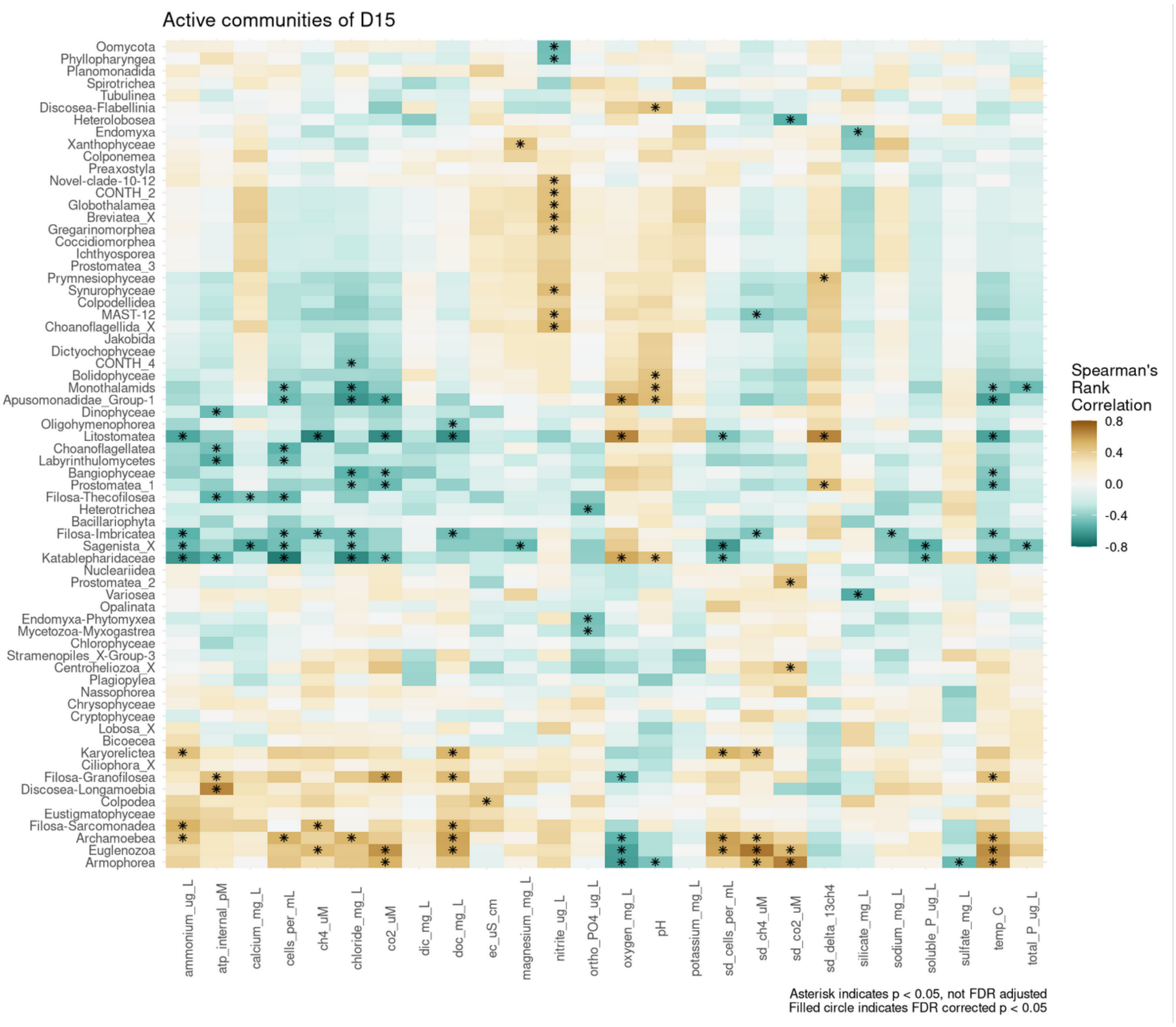


**Figure S4.** Spearman's rank correlations between protistan classes and environmental variables for the active community of D15 groundwater monitoring site. The heatmap is colored based on the correlation strength, with blue shades for negative correlations and brown shades for positive correlations. Significant correlations are marked with an asterisk (uncorrected).

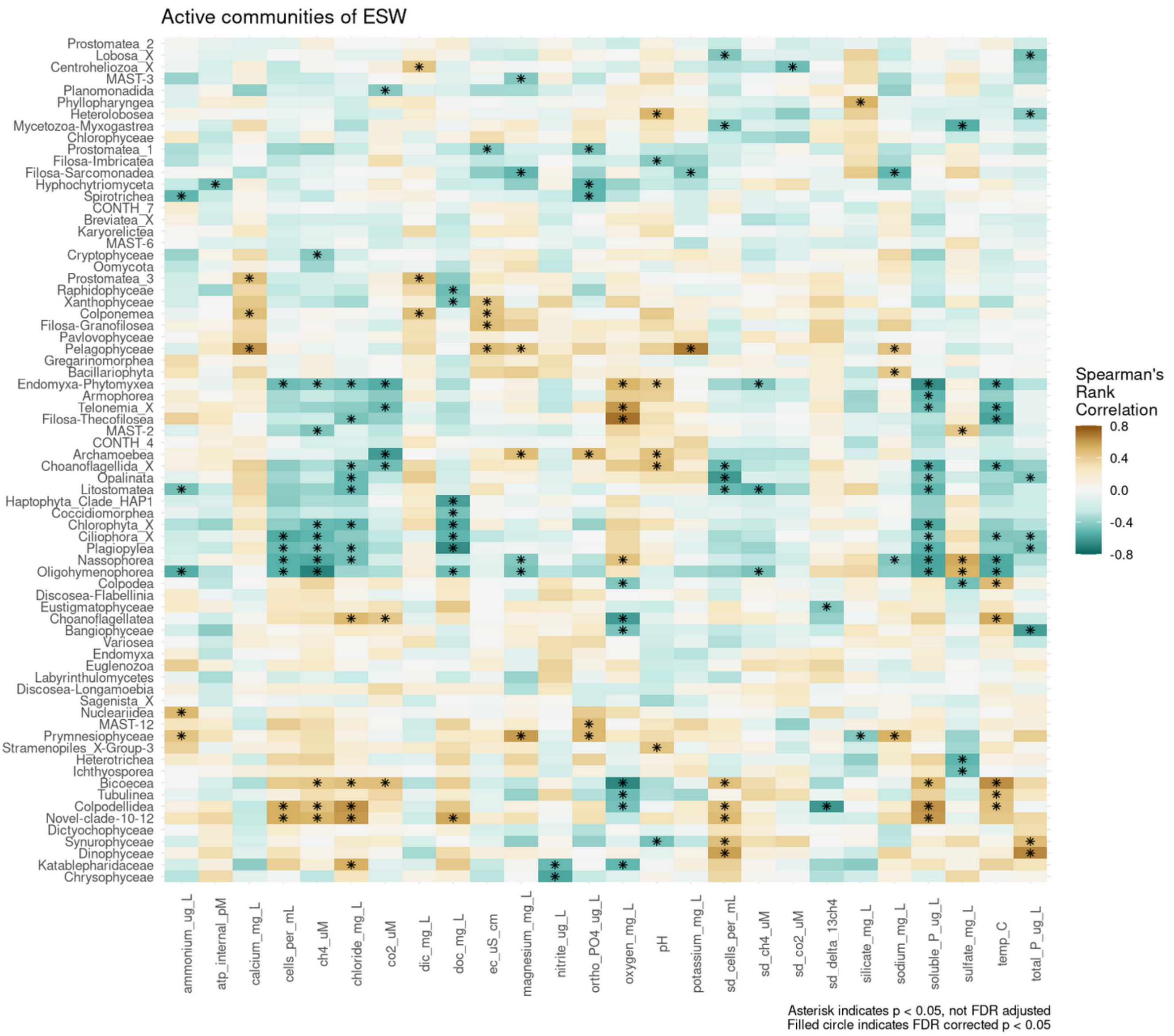


**Figure S5.** Spearman's rank correlations between protistan classes and environmental variables for the active community of ESW surface water site. The heatmap is colored based on the correlation strength, with blue shades for negative correlations and brown shades for positive correlations. Significant correlations are marked with an asterisk (uncorrected).
